# An abscisic acid (ABA) homeostasis regulated by its production, catabolism and transport in peanut leaves in response to drought stress

**DOI:** 10.1101/569848

**Authors:** Haitao Long, Zhao Zheng, Yajun Zhang, Pengzhan Xing, Xiaorong Wan, Yixiong Zheng, Ling Li

**Affiliations:** School of Life Sciences, South China Normal University, Guangzhou, China; College of Agriculture and Biology, Zhongkai University of Agriculture and Engineering, Guangzhou, China

**Author notes:** Corresponding authors (XW); (LL). **Data Availability:** All relevant data are within the manuscript.

**Keywords:** Peanut (*Arachis hypogaea* L.), leaves, drought stress, ABA homeostasis, production and catabolism, transport

## Abstract

ABA is an important messenger that acts as the signaling mediator for regulating the adaptive response of plants to drought stress. Two production pathways, *de novo* biosynthesis and hydrolysis of glucose-conjugated ABA by β-glucosidase (BG), increase cellular ABA levels in plants. ABA catabolism via hydroxylation by 8’-hydroxylase (CYP707A), or conjugation by uridine diphosphate glucosyltransferase (UGT), decreases cellular ABA levels. The transport of ABA through ATP-binding cassette (ABC)-containing transporter proteins, members of ABC transporter G family (ABCG), across plasma membrane (PM) is another important pathway to regulate cellular ABA levels. In this study, based on our previously constructed transcriptome of peanut leaves in response to drought stress, fourteen candidate genes involved in ABA production (including *AhZEP*, *AhNCED1* and *AhNCED3*, *AhABA2*, *AhAAO1* and *AhAAO2*, *AhABA3*, *AhBG11* and *AhBG24*), catabolism (including *AhCYP707A3*, *AhUGT71K1* and *AhUGT73B4*) and transport (including *AhABCG22-1* and *AhABCG22-2*), were identified homologously and phylogenetically, and further analyzed at the transcriptional level by real-time RT-PCR, simultaneously determining ABA levels in peanut leaves in response to drought. The high sequence identity and very similar subcellular localization of the proteins deduced from 14 identified genes involved in ABA production, catabolism and transport with the reported corresponding enzymes in databases suggest their similar roles in regulating cellular ABA levels. In response to drought stress, ABA accumulation levels in peanut leaves agree very well with the up-regulated expressions of ABA-producing genes (*AhZEP*, *AhNCED1*, *AhAAO2*, *AhABA3*, *AhBG11* and *AhBG24*) and PM-localized ABA importer genes (*AhABCG22-1* and *AhABCG22-2*), although the expression of ABA catabolic genes (*AhCYP707A3* and *AhUGT71K1*) was also up-regulated. It is likely that drought-responsive induction of catabolic genes helps not only to maintain ABA levels within a permissible range, but also to prepare the plant for degradation of ABA after removal of the stress. These results suggest that ABA homeostasis in peanut leaves in response to drought may be coordinated by a master regulatory circuit that involves production, catabolism, and as well as transport.

## Introduction

The plant hormone ABA plays pivotal roles in many important physiological processes including stomatal closure, seed dormancy, growth and various abiotic stress responses [1,2]. ABA is mainly produced by the *de novo* biosynthetic pathway through the oxidative cleavage of carotenoids [3]. In this pathway, zeaxanthin epoxidase (ZEP/ABA1) catalyzes the formation of all transviolaxthin from zeaxanthin [4]. Nine *cis*-epoxycarotenoid dioxygenase (NCED) cleaves carotenoids to form xanthoxin [5,6]. Xanthoxin is assumed to be transported from the plastids to the cytosol, although the precise mechanism that mediates this transport is not yet known [2]. The short-chain alcohol dehydrogenase/reductase (SDR/ABA2) converts xanthoxin derived from cleavage of carotenoids into abscisic aldehyde [7,8], which is finally oxidized into ABA by abscisic aldehyde oxidase (AAO) [9–11]. Aldehyde oxidase requires the molybdenum cofactor sulfurase/ABA3 to produce a functional cofactor for its catalytic activity [12]. All of the steps of ABA *de novo* biosynthesis occur in plastids except for the final two stages, which take place in the cytosol [9–11].

An alternative pathway for producing ABA is via hydrolysis of the ABA-glucosyl ester (ABA-GE), which is an inactive glucose-conjugated form of ABA. Intracellular ABA-GE can be hydrolysed by the two β-glucosidase (BG) homologs AtBG1 and AtBG2 in *Arabidopsis* [13,14], which localize to the endoplasmic reticulum (ER) and vacuole, respectively. The single-step reaction of β-glucosidase-regulated hydrolysis of ABA-GE to ABA is an ideal and important way to achieve the rapid increase in ABA levels necessary for plants to meet their physiological needs [14].

ABA catabolism is also a mechanism for regulating ABA levels. In *Arabidopsis* it proceeds mainly via two pathways, namely ABA 8’-hydroxylation catalyzed by ABA 8’-hydroxylase, the cytochrome P450 (CYP) 707A family [15], and ABA conjugation with glucose mediated by glucosyltransferases [16,17]. The 8′-hydroxylation of ABA is mediated by CYP707A family of proteins (CYP707As 1, 2, 3 and 4) in *Arabidopsis* [15]. We previously reported two genes (*AhCYP707A1* and *AhCYP707A2*) encoding ABA 8’-hydroxylase from peanut [18]. Expressions of *AhCYP707A1* and *AhCYP707A2* genes were ubiquitous in peanut roots, stems and leaves with different transcript levels, and were regulated osmotically, as shown by responses to osmotic stress instead of ionic stress [18]. The different spatial and temporal patterns of expression of four *Arabidopsis* and two peanut *CYP707A* genes, suggesting that each of the gene products may function in different physiological or developmental processes. The expression of all four Arabidopsis *CYP707A* genes was induced by dehydration stress and subsequent rehydration [15,19], which indicates that ABA levels are regulated by a balance between biosynthesis and catabolism, including feedback-induced catabolism. Conjugation of ABA with glucose is catalysed by ABA-uridine diphosphate (UDP) glucosyltransferases (UGTs), which include *Arabidopsis* UGT71B6 and its two closely related homologs, UGT71B7 and UGT71B8 [16,17]. A recent study has shown that UGT71B6, UGT71B7 and UGT71B8 play crucial roles in ABA homeostasis and adaptation to dehydration, osmotic and high-salinity stresses in *Arabidopsis* [17]. ABA catabolic pathways appear to be localized in the cytosol (UGT71Bs) and the ER membrane (CYP707As) [20].

Moreover, ABA and its metabolites are transported between subcellular compartments within a cell as well as between cells [2,20]. For the regulation of endogenous ABA level in plants, it is still crucial to determine how ABA transport is regulated, and whether it is involved in the control of physiological responses. The protonated form of ABA can easily cross biological membranes because of its hydrophobicity. So ABA could be transported from relatively low-pH to high-pH cellular compartments via a passive diffusion that does not require specific transporters [21]. The first step in ABA transport might be ABA export out of cells. ABA is synthesized in the cytosol, where the pH is relative higher than that in the apoplastic space. Therefore a specific transporter may be required for ABA export to the apoplastic space. Recent studies in *Arabidopsis* have identified both ABA exporters and ABA importers localized to the plasma membrane (PM). ABA transporters were first identified in *Arabidopsis*, and they are ATP-binding cassette (ABC)-containing transporter proteins, members of ABC transporter G family [22,23]. *AtABCG25* encodes a half-size ABC transporter protein and which is responsible for exporting ABA from vascular tissues, the main sites of ABA synthesis in plants [22]; AtABCG40, a full-size ABC transporter, acts as an ABA importer in plant cells [23]. Subsequently, AtABCG22 was shown to be required for stomatal regulation and proposed to function as ABA transporter [24]. Currently, Kang et al [25] have demonstrated that four AtABCG proteins function together to supply ABA in mature imbibed seeds of *Arabidopsis*. They showed that AtABCG25 and AtABCG31 export ABA from the endosperm to the embryo, whereas AtABCG30 and AtABCG40, transport ABA into the embryo [25]. The low-affinity nitrate transporter (NRT1) was also reported to function as an ABA importing transporter (AIT1) [26,27]. Zhang et al [28] showed that AtDTX50 (Detoxification Efflux Carrier 50), a membrane protein in the MATE (Multidrug and Toxic Compound Extrusion) transporter family in *Arabidopsis*, mediated ABA efflux from the cytosol of vascular and guard cells. Recently, we have also isolated an ABA transporter-like 1 gene (*AhATL1*) from peanut plants, which modulated ABA sensitivity through specifically affecting ABA import into cells in transgenic *Arabidopsis* [29]. It appears that multiple types of transporters are involved in ABA transport in plants. Therefore, ABA-specific transporters localized to the plasma membrane also regulate the cellular ABA levels in plant cells.

Drought is one of the major abiotic stresses that limit the growth and production of plants. The mechanisms of drought stress response have been investigated most extensively in *Arabidopsis*, which include ABA-dependent and non-ABA dependent pathways [1,30,31]; ABA homoeostasis modulated by its production, inactivation, and transport is considered to play vital roles for plant development and stress responses; the transcriptional regulation of genes involved in either ABA production or ABA inactivation is of great importance in ABA homoeostasis [32]. However, our knowledge of the genes involved in regulation of ABA levels is relatively rare in agricultural crops in response to drought. We have used peanut, an economically important oil and protein rich crop, to address the issue [18,33–42]. In the present study, based on the screening of our previously constructed transcriptome of peanut leaves in response to drought stress [38], we report the identification and expression analysis of genes encoding the enzymes involved in ABA production [including one *ZEP* (*AhZEP*), two *NCED*s (*AhNCED1* and *AhNCED3*), one *ABA2* (*AhABA2*), two *AAO*s (*AhAAO1* and *AhAAO2*), one *ABA3* (*AhABA3*), and two *BG*s (*AhBG11* and *AhBG24*)], catabolism [including one *CYP707A* (*AhCYP707A3*) and two *UGT*s (*AhUGT71K1* and *AhUGT73B4*)], and transport [including two *ABCG*s (*AhABCG22-1* and *AhABCG22-2*), which jointly contribute to the regulation of ABA homeostasis precisely in peanut leaves in response to drought.

## Materials and methods

### Plants and growth conditions

Seeds of peanut (*Arachis hypogaea* L. cv ‘Yueyou 7’) were sown in pots with a potting mixture of vermiculite, perlite and soil (1:1:1), and grown in a growth chamber with 16 h of light from fluorescent and incandescent lamps (200 μmol m^−2^ s^−2^) followed by 8 h of darkness at 28°C. Plants were watered daily with half-strength Murashige and Skoog nutrient solution [43].

### Drought stress treatment of plants

For the treatment of polyethylene glycol (PEG6000)-simulated drought stress, three-leaf-stage (10-15 days after planting) peanut plants were removed from the soil mixture carefully to avoid injury, and then hydroponically grown in a solution for indicated time containing 20% (W/V) PEG6000, or deionized water as a control, respectively. For all these treatments, peanut leaves were frozen in liquid nitrogen immediately following the treatments and stored at −80°C until analysis. The entire experiments were biologically repeated at least three times.

### Molecular cloning of genes encoding enzymes involved in ABA biosynthesis, catabolism and transport from peanut

From the constructed transcriptome which contained 47 842 assembled unigenes of three-leaf-stage peanut leaves in response to drought [38], we screened the fragments of genes encoding the enzymes involved in ABA production (including *AhZEP*, *AhNCED1* and *AhNCED3*, *AhABA2*, *AhAAO1* and *AhAAO2*, *AhABA3*, *AhBG11* and *AhBG24*), catabolism (including *AhCYP707A3*, *AhUGT71K1* and *AhUGT73B4*) and transport (including *AhABCG22-1* and *AhABCG22-2*). The missing 5’ and 3’ ends of the screened genes were obtained by rapid amplification of cDNA ends (RACE) using the GeneRacer kit according to the manufacturer’s instructions (ThermoFisher, Shanghai, China). The gene specific primers for 5’ and 3’ RACE of target genes were listed in Table 1. In all cloning experiments, PCR fragments were gel-purified and ligated into the pMD 19-T Vector (TaKaRa, Dalian, China), and confirmed by sequencing from both strands.

### Sequence analyses and alignments

The routine sequence analyses were performed by using the Gene Runner (Hastings Software, Inc., New York, USA). Computer analyses of cDNAs and deduced amino acid sequences were carried out using the Basic Local Alignment Search Tool (BLAST) program at the National Center for Biotechnology Information Services [44]. Multiple alignments of deduced amino acid sequences from target genes were performed by using the Clustal W program in the BioEdit software (Isis Pharmaceuticals, Inc., Carlsbad, USA). The full-length protein sequences were phylogenetically analyzed by using the MEGA 4 software with a bootstrapping set of 1000 replicates [45]. The subcellular localization of target proteins was predicted by using the iPSORT algorithm [46] at the website: http://ipsort.hgc.jp/ and the WoLF PSORT tool at the website: http://www.genscript.com/wolf-psort.html.

**Table 1.**
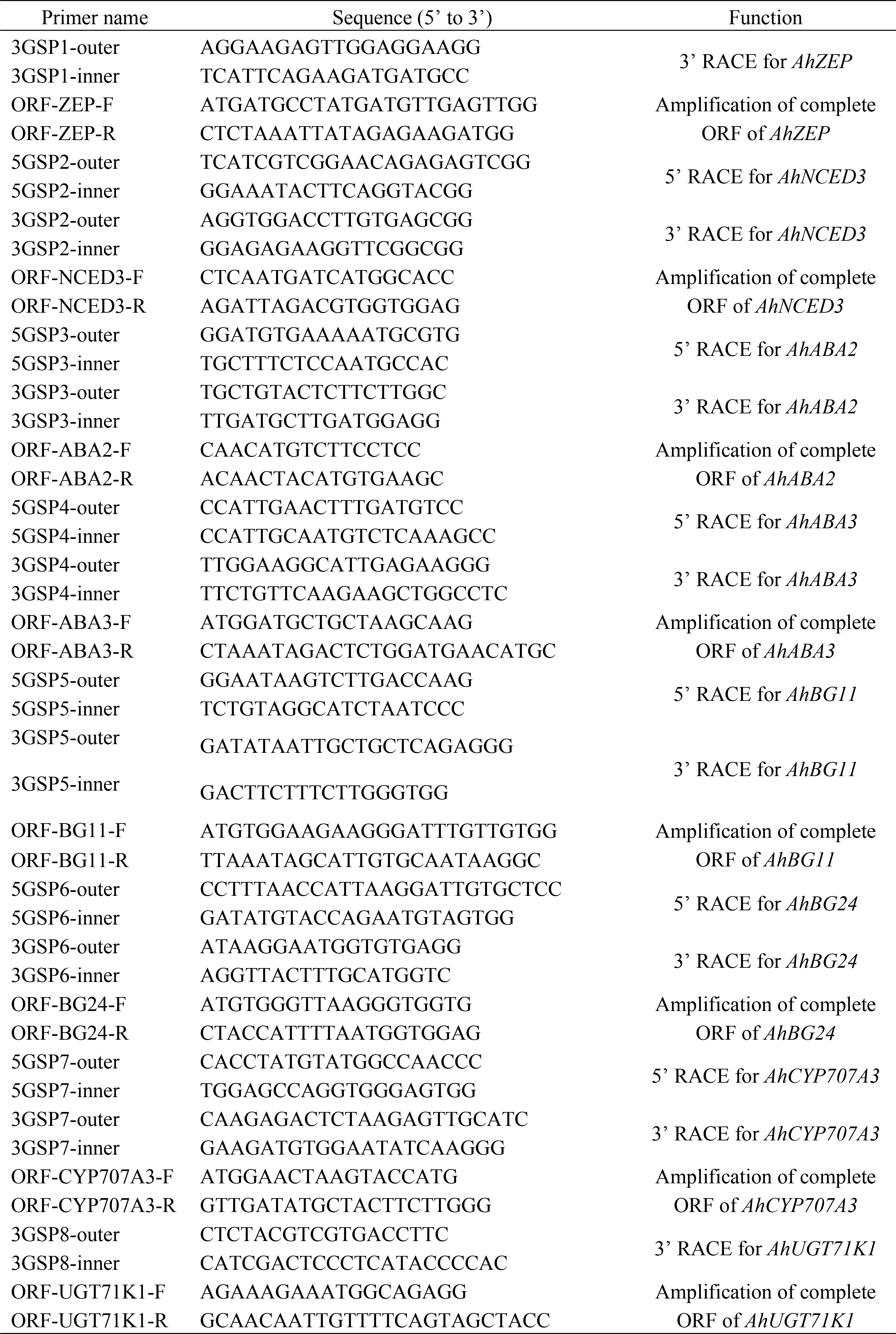
Primer sequences used in the present study.

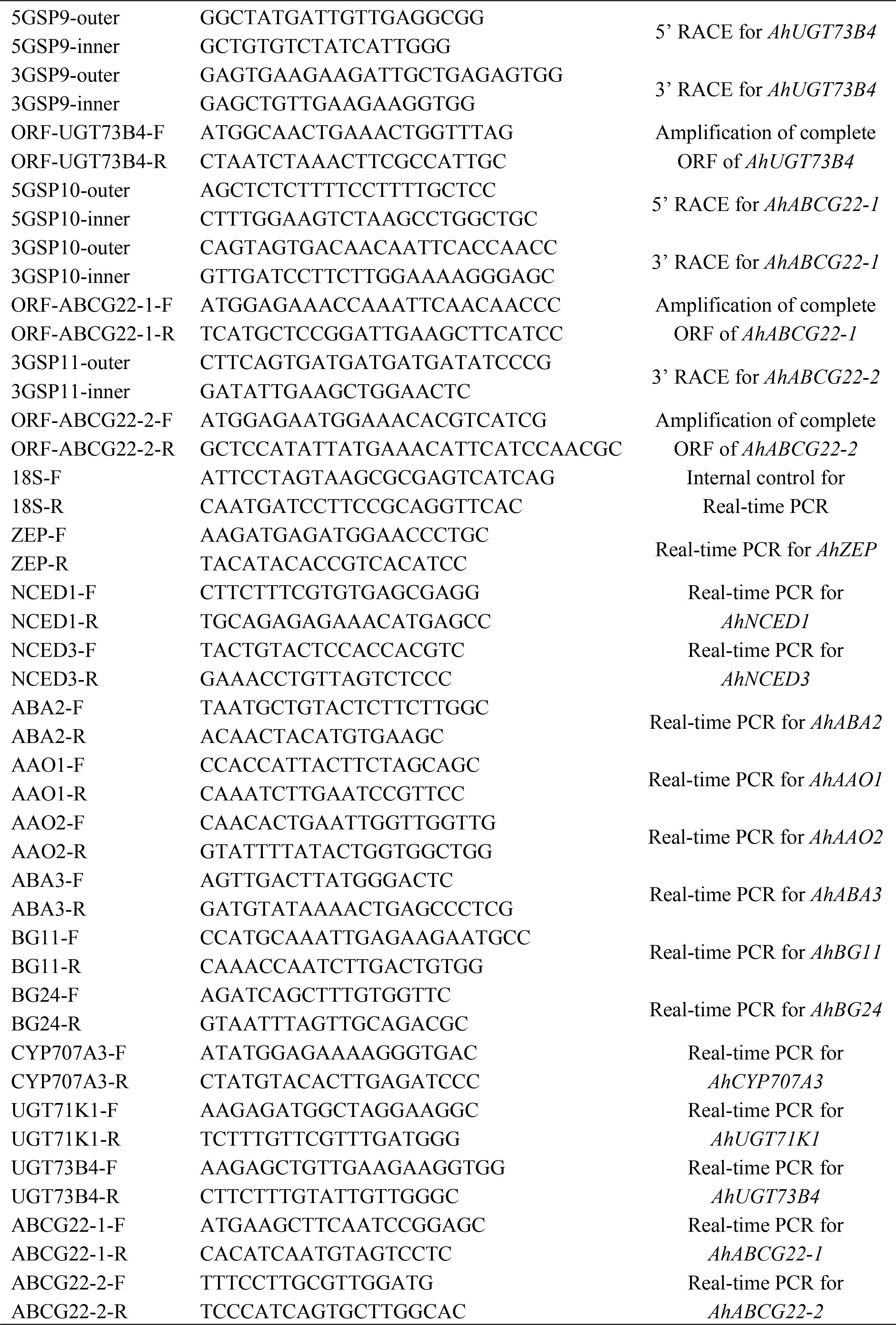

### Real-time quantitative RT-PCR performance

The isolated RNA by using the modified phenol chloroform method as previously described [33] was treated with RNase-free DNase I (TaKaRa, Dalian, China) at 37°C for 1 h to eliminate DNA contamination in real-time quantitative RT-PCR analysis. Two micrograms of total RNA and 200 ng of a random primer were used in reverse transcription (RT) through the cDNA synthesis kit (TaKaRa, Dalian, China) according to the manufacturer’s protocol. To investigate the expressions of target genes in peanut leaves in response to drought, the gene-specific primers were designed and listed in Table 1 to amplify the each corresponding cDNA for real-time quantitative PCR. As an internal control for normalization of target gene expression, the primers 18S-F (5’-ATT CCT AGT AAG CGC GAG TCA TCA G-3’) and 18S-R (5’-CAA TGA TCC TTC CGC AGG TTC AC-3’) specific to peanut 18S rRNA gene (GenBank accession no. AF156675) were used to amplify a fragment of 226 bp. Real-time quantitative PCRs were performed in the presence of Power SYBR green PCR Master Mix (Applied Biosystems, Guangzhou, China). Amplification was monitored in real-time with the MiniOpticon™ Real-Time PCR System (Bio-Rad, Shanghai, China). The products of real-time quantitative PCR were confirmed by determining the melt curves for the products at the end of each run, by analysis of the products using gel electrophoresis, and by sequencing. Quantification of the normalized gene expression was performed with the comparative cycle threshold (Ct) method [47]. Three biological and three technical replicates were performed for each experiment. All RT-PCR data were expressed as the mean ± standard error. Statistical differences of expressions of target genes were assessed by one-way analysis of variance (ANOVA) followed by the least significant difference (LSD) and Student-Neumann-Keuls (SNK) post hoc comparison. The analyses were performed with SPSS 13.0 software (SPSS Inc., Chicago, IL, USA). The threshold of significance was defined as p<0.05.

### Measurement of endogenous ABA level

Endogenous ABA was extracted from the frozen leaf sample as described by Xiong et al [12]. Extraction in non-oxidative methanol:water (80/20, v/v), pre-purification through SepPak C 18 cartridges (Waters, Milford, MA, USA), and HPLC fractionation in a Nucleosil C 18 column (Mecherey-Nagel, Germany) have been previously described [39,48]. The ELISA procedure was performed based on the competition, for a limited amount of monoclonal anti-ABA antibody (1:2000 in 5% BSA/PBS), between standard ABA-BSA conjugate absorbed on the wells of a microtitration plate and free ABA extracted from the samples. Bound antibodies were labeled with a peroxidase-conjugated goat antibody to mouse immunoglobulins (Sigma), and peroxidase activities were then determined. A standard carve was established for each microtitration plate. The ABA level was determined triplicately with three replicates for each.

## Results and discussion

### Characterization of genes encoding enzymes involved in ABA production, catabolism and transport from peanut

From the constructed transcriptome of three-leaf-stage peanut leaves in response to drought [38], fourteen candidate genes involved in ABA production (*AhZEP*, *AhNCED1* and *AhNCED3*, *AhABA2*, *AhAAO1* and *AhAAO2*, *AhABA3*, *AhBG11* and *AhBG24*), catabolism (*AhCYP707A3*, *AhUGT71K1* and *AhUGT73B4*) and transport (*AhABCG22-1* and *AhABCG22-2*), were screened and identified homologously and phylogenetically. The characteristics of the full-length cDNAs of fourteen screened target genes obtained by RACE and the corresponding deduced proteins were shown in Table 2.

**Table 2.** Characteristics of full-length cDNAs of fourteen screened target genes and corresponding deduced proteins.

| Gene | Accession no. | cDNA full-length (bp) | Open read frame (ORF) (bp) | 5' Untranslated region (UTR) (bp) | 3' UTR (bp) | Amino acid residues of deduced protein | Molecular weight of deduced protein (kDa) | Isoelectric point of deduced protein | Subcellular localization of deduced protein |
| --- | --- | --- | --- | --- | --- | --- | --- | --- | --- |
| *AhZEP* | MH037805 | 495 | 270 | 111 | 111 | 90 | 10.15 | 4.06 | Chloroplast |
| *AhNCED1* | AJ574819 | 2486 | 1803 | 187 | 493 | 601 | 66.86 | 8.39 | Chloroplast |
| *AhNCED3* | MH037806 | 2263 | 1917 | 181 | 162 | 639 | 70.88 | 6.91 | Chloroplast |
| *AhABA2* | MH037807 | 1082 | 852 | 130 | 97 | 284 | 30.23 | 6.60 | Cytosol |
| *AhAAO1* | EU183360 | 4585 | 3801 | 326 | 455 | 1267 | 138.04 | 6.38 | Cytosol |
| *AhAAO2* | EU816196 | 4677 | 4080 | 327 | 267 | 1360 | 149.41 | 6.55 | Cytosol |
| *AhABA3* | MH037808 | 3719 | 2448 | 112 | 1156 | 816 | 91.0 | 6.93 | Cytosol |
| *AhBG11* | MH037809 | 1970 | 1539 | 228 | 200 | 513 | 58.84 | 5.68 | Endoplasmic reticulum (ER) |
| *AhBG24* | MH037810 | 1763 | 1488 | 61 | 211 | 496 | 55.86 | 8.40 | Vacuolar |
| *AhCYP707A3* | MH037811 | 1930 | 1392 | 163 | 372 | 464 | 53.15 | 9.29 | ER |
| *AhUGT71K1* | MH037812 | 1628 | 1440 | 54 | 131 | 480 | 53.62 | 5.65 | Cytosol |
| *AhUGT73B4* | MH037813 | 1607 | 1452 | 87 | 65 | 484 | 54.41 | 5.42 | Cytosol |
| *AhABCG22-1* | MH037814 | 2572 | 2253 | 154 | 162 | 751 | 83.10 | 8.98 | Plasma membrane |
| *AhABCG22-2* | MH037815 | 2429 | 2193 | 66 | 167 | 731 | 81.12 | 7.89 | Plasma membrane |

The main pathways of *de novo* ABA biosynthesis occur both in plastids and in the cytosol, starting from the precursor isopentenyl diphosphate (IPP), which is synthesized primarily in plastids from glyceraldehyde 3-phosphate and pyruvate, resulting in the successive production of the intermediates phytoene and lycopene [2,3]. Cyclization and hydroxylation of lycopene produce the oxygenated carotenoid zeaxanthin, which is then catalyzed by zeaxanthin epoxidase (ZEP) encoded by the *Arabidopsis AtABA1* locus to synthesize the violaxanthin [48]. In the present study, from our constructed drought-induced transcriptome of peanut leaves, one candidate *ZEP* was identified as *AhZEP*, encoding the enzyme AhZEP which shared 81%, 79%, 78%, 73% and 73% sequence identity with *Glycine soja* GsZEP (KHN42080), *Vigna radiata* VrZEP1 (XP_022631763), *Medicago truncatula* MtZEP (XP_013453497), *Medicago sativa* (AIP98334), and *Lupinus luteus* LlZEP (AHI87686), respectively. AhZEP protein was predicted by the iPSORT algorithm to have a chloroplast transit peptide MMPMMLSWVLGGNSSKLEGRPVCCRLSDKA at the N-terminus.

In *Arabidopsis*, five AtNCEDs (AtNCED2, 3, 5, 6 and 9) were characterized to cleave the substrates violaxanthin and neoxanthin to a C_15_ product, xanthoxin (the first cytoplasmic precursor in ABA biosynthetic pathway) [49]. Here two candidate *NCED* genes, *AhNCED1* (our previous work [33,34]) and *AhNCED3* were characterized from the constructed drought-induced transcriptome of peanut leaves. Multiple alignments showed that the deduced amino acids from *AhNCED1* and *AhNCED3* shared 59.2% sequence identity with each other. AhNCED3 protein shared 60.2%, 62.2%, 61.9%, 47.9% and 54.7% sequence identity with *Arabidopsis* AtNCED2, 3, 5, 6 and 9, respectively. A putative 30-amino-acid chloroplast transit peptide MIMAPSSIALNSASSSTWAKKPHQLSRPFS predicted by the iPSORT algorithm is located at the N-terminus of AhNCED3 protein, structurally similar with all reported NCED proteins [3,33,49,50]. Phylogenetic analysis of AhNCED1, AhNCED3 and five *Arabidopsis* NCEDs showed that AhNCED1 and AtNCED3 were clustered into one group (Fig 1), both of them playing a vital role in stress-induced ABA biosynthesis in leaves [34,49]. AhNCED3 was clustered with AtNCED2 and AtNCED5 (Fig 1), which accounted for the main *NCED* transcripts in flowers [49].

**Fig 1.**
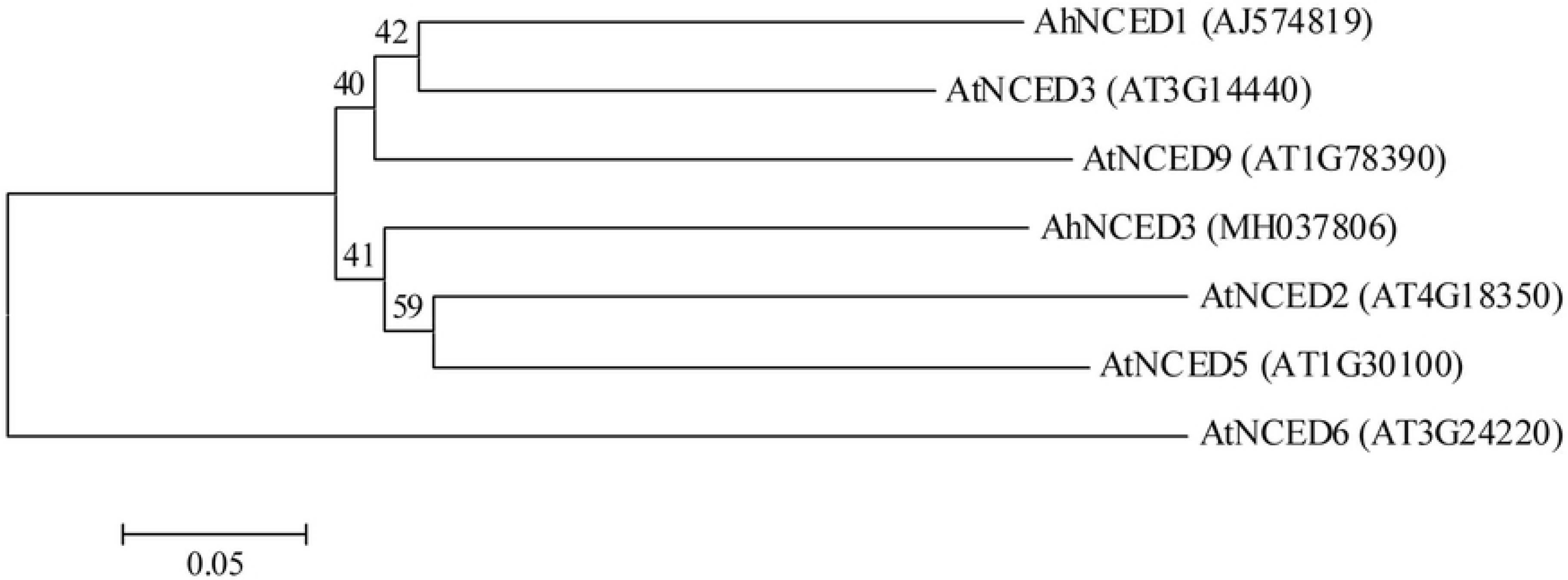
Phylogenetic analysis of amino acid sequences deduced from *AhNCED1*, *AhNCED3*, and five *Arabidopsis NCEDs* (*AtNCED2*, *3*, *5*, *6* and *9*) Multiple sequence alignment was performed using Clustal W and phylogenetic tree was constructed via the Neighbor-Joining method in MEGA 4 software. Bootstrap values from 1000 replicates for each branch were shown. GenBank accession numbers for each aligned NCED sequence were indicated in parentheses. The scale bar is 0.05.

The conversion of xanthoxin into abscisic aldehyde is catalyzed by AtABA2 in *Arabidopsis*, which belongs to the short-chain dehydrogenases/reductases (SDR) family [7,8]. A severe ABA deficiency resulting from loss of function of *AtABA2* suggests that AtABA2 protein appears to be encoded by a single gene in *Arabidopsis* genome [8]. In the present study, *AhABA2* was characterized to encode AtABA2 homolog in peanut. Multiple alignments showed that AhABA2 protein shared 67.2%, 70% and 67.9 sequence identity with AtABA2, tomato SlABA2 and tobacco NtABA2, respectively (Fig. 2A). The domain (residues 3 to 285 in AtABA2) with xanthoxin dehydrogenase activity was highly conserved in all aligned ABA2 proteins (Fig 2A). AhABA2 was phylogenetically closer to soybean GmABA2 in the leguminous cluster (Fig 2B).

**Fig 2.**
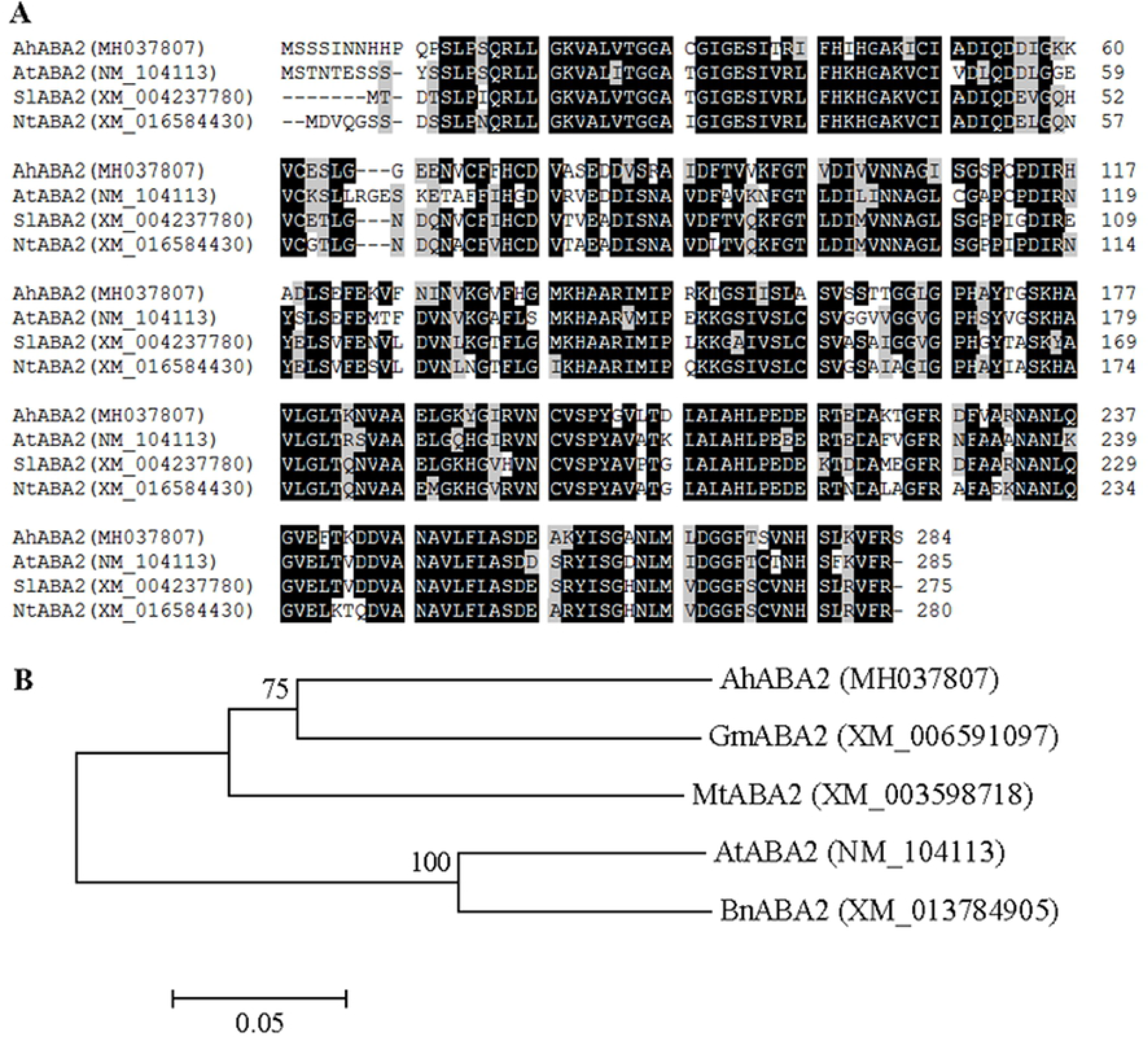
Sequence analyses of ABA2 proteins from peanut, *Arabidopsis*, tomato, tobacco, soybean, alfalfa, and winter rape. **(A)** Alignment of deduced amino acid sequences from peanut *AhABA2*, *Arabidopsis AtABA2*, tomato *SlABA2*, and tobacco *NtABA*2. Identical and similar amino acid residues were shaded in black and gray, respectively. Dotted lines indicated gaps that were introduced to maximize the alignment. Amino acids were numbered from the initial methionine. GenBank accession numbers for each aligned ABA2 homolog were indicated in parentheses. **(B)** Phylogenetic analysis of amino acid sequences of AhABA2, AtABA2, soybean GmABA2, alfalfa MtABA2, and winter rape BnABA2. Multiple sequence alignment was performed using Clustal W and phylogenetic tree was constructed via the Neighbor-Joining method in MEGA 4 software. Bootstrap values from 1000 replicates for each branch were shown. GenBank accession numbers for each analyzed ABA2 were indicated in parentheses. The scale bar is 0.05.

The oxidation of abscisic aldehyde to ABA, which is catalyzed by abscisic aldehyde oxidase, is the final step in ABA biosynthetic pathway. Among four abscisic aldehyde oxidases (AtAAO1 to 4) in *Arabidopsis*, AtAAO3 was reported to actively utilize abscisic aldehyde as a substrate, most probably the only one AAO involved in ABA biosynthesis [11]. Here our previously characterized two peanut *AAO* genes, *AhAAO1* [51] and *AhAAO2* [52], were also screened from the constructed drought-induced transcriptome of peanut leaves. AhAAO1 protein was predicted to localize in the cytosol by the WoLF PSORT tool, and AhAAO2 was predicted by the iPSORT algorithm as not having any of signal, mitochondrial targeting, or chloroplast transit peptides. The aldehyde oxidase requires a molybdenum cofactor (MoCo) for its catalytic activity. To date, *AtABA3* (a single-copy gene in the genome) was the only reported *ABA3* gene encoding *Arabidopsis* sulfurase that produces a functional cofactor [12]. In this study, an *AtABA3* homolog gene *AhABA3* was characterized from the drought-induced transcriptome of peanut leaves. Multiple alignments showed that AhABA3 protein shared 82.1%, 80.9% and 61.3% sequence identity with soybean GmABA3, *Cajanus cajan* CcABA3 and AtABA3, respectively (Fig 3). The putative pyridoxal phosphate (PLP) binding motif and the conserved cysteine motif identified by Xiong et al [12] both exist in AhABA3 protein sequence (Fig 3).

**Fig 3.**
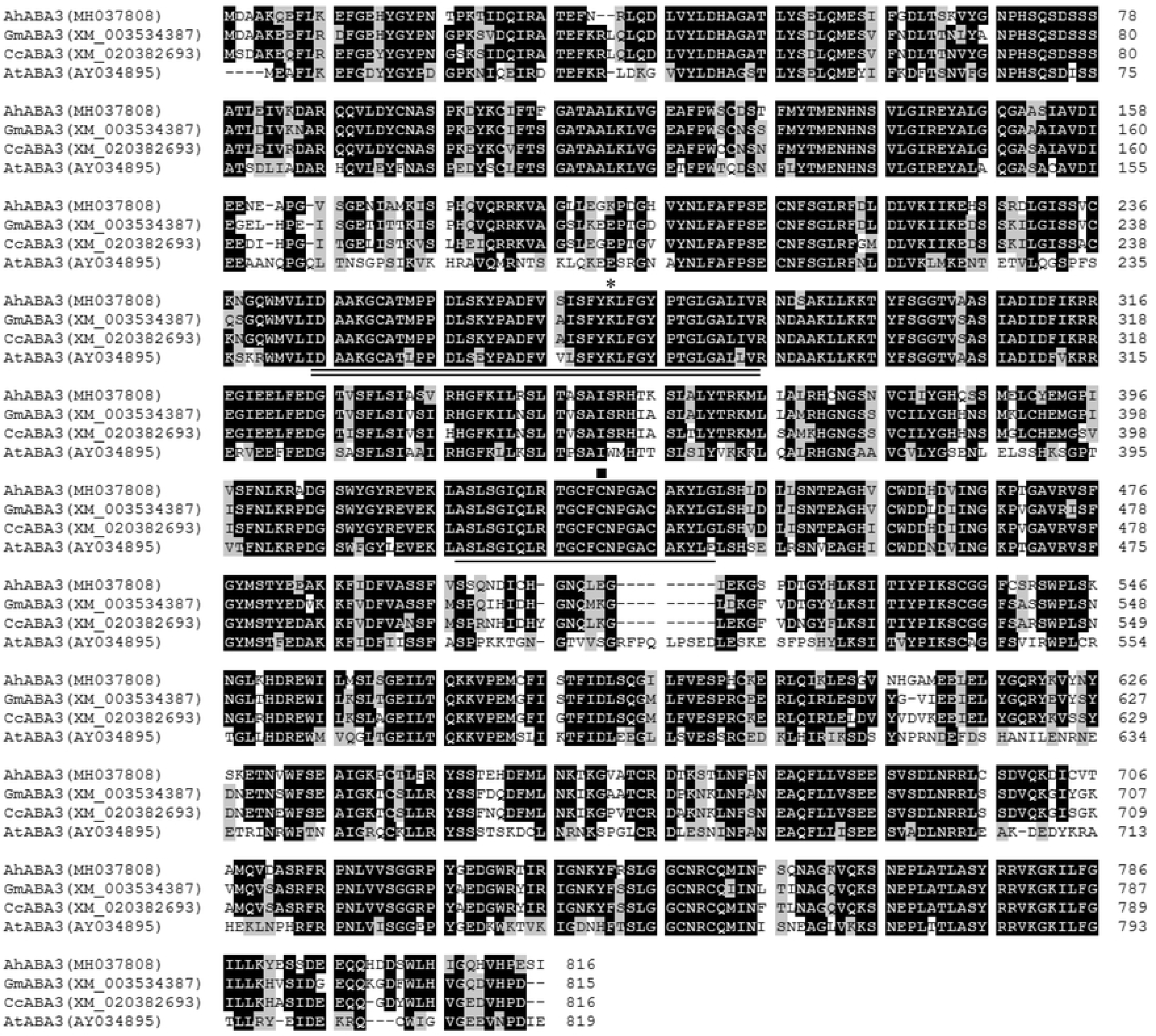
Alignment of deduced amino acid sequences from peanut *AhABA3*, soybean *GmABA3*, *Cajanus cajan CcABA3* and *Arabidopsis AtABA3*. Identical and similar amino acid residues were shaded in black and gray, respectively. Dotted lines indicated gaps that were introduced to maximize the alignment. The conserved cysteine motif was underlined and the putative PLP binding motif was double underlined. The conserved critical lysine residue in the PLP domain was indicated with an asterisk, and the conserved cysteine residue was indicated with a square. Amino acids were numbered from the initial methionine. GenBank accession numbers for each aligned ABA3 homolog were indicated in parentheses.

The hydrolysis of ABA-GE catalyzed by β-glucosidase (BG) is an alternative pathway to produce ABA. The β-glucosidase homologs, *Arabidopsis* AtBG1 and AtBG2, localize to the ER and vacuole, respectively [13,14]. AtBG2 belongs to the same subfamily as AtBG1 that consists of 16 members in the large number of β-glucosidases found in *Arabidopsis* [13,53], which can be divided into two groups: AtBG1 belongs to the group of seven members with an ER retrieval signal, and AtBG2 belongs to the other group of nine members without the ER retrieval signal [14]. In the present study, two *BG* homologs, *AhBG11* and *AhBG24*, were characterized from our constructed drought-induced transcriptome of peanut leaves. AhBG11 protein shared 41.6%, 37.7% and 32.5% sequence identity with AhBG24, AtBG1 and AtBG2, respectively; and AhBG24 shared 40.2% and 37.2% sequence identity with AtBG1 and AtBG2, respectively (Fig 4). AhBG11 and AhBG24 were predicted by the WoLF PSORT tool to localize to the ER and vacuole, respectively (Table 2; Fig 4), suggesting that AhBG11 and AhBG24 might belong to the group with AtBG1 and the other group with AtBG2, respectively.

**Fig 4.**
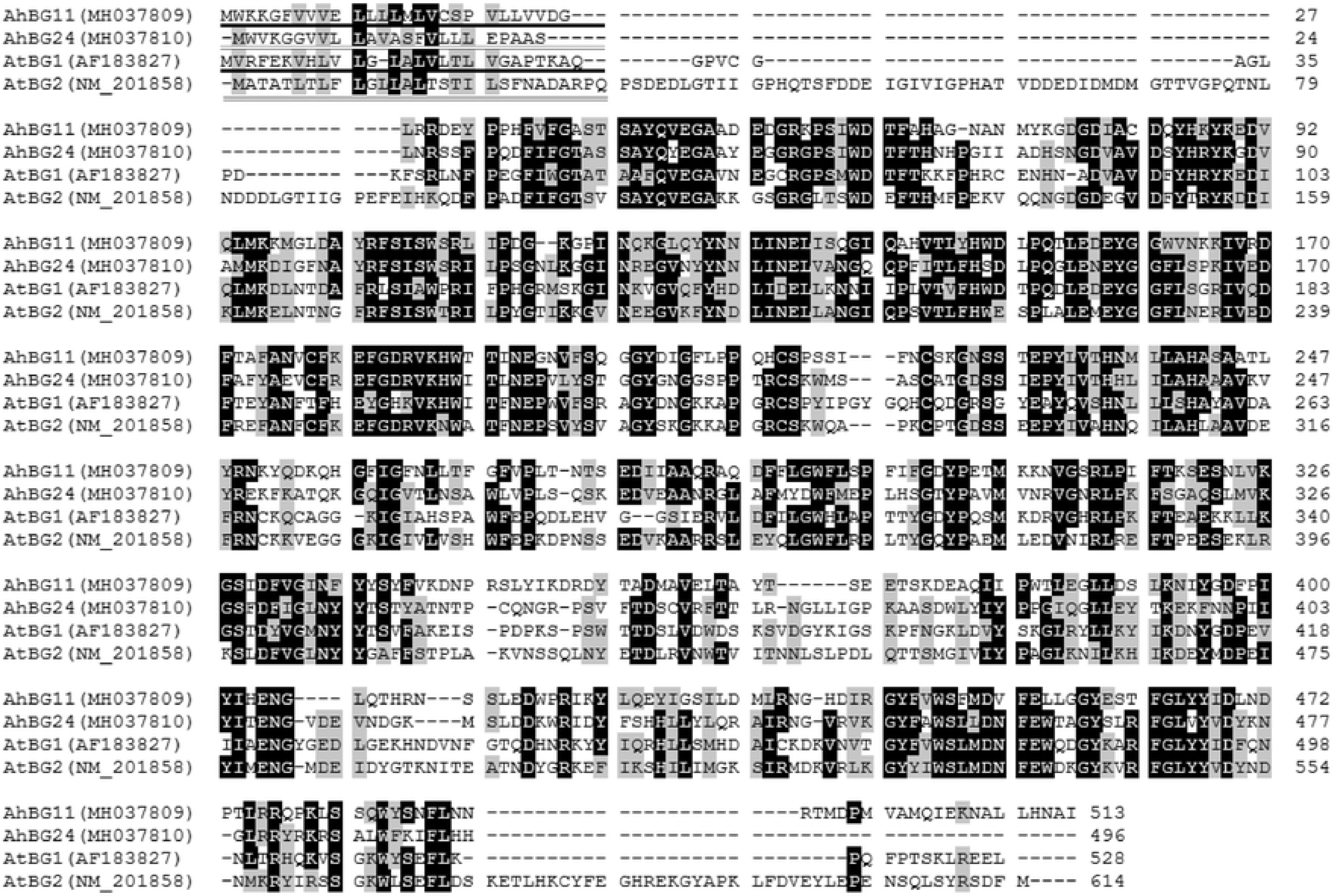
Alignment of deduced amino acid sequences from peanut *AhBG11*, *AhBG24*, and *Arabidopsis AtBG1*, *AtBG2*. Identical and similar amino acid residues were shaded in black and gray, respectively. Dotted lines indicated gaps that were introduced to maximize the alignment. Putative ER-localization signal peptide was underlined in AtBG1 and AhBG11 [13]; and putative vacuolar-targeting motif was double underlined in AtBG2 and AhBG24 [14]. Amino acids were numbered from the initial methionine. GenBank accession numbers for each aligned BG homolog were indicated in parentheses.

The catabolic process of ABA mainly involves two pathways, hydroxylation and glucose conjugation. The 8′-hydroxylation of ABA is the predominant enzymatic reaction, which is mediated by the protein encoded by AtCYP707A gene family (*AtCYP707A1*, *2*, *3* and *4*) in *Arabidopsis* [15]. In this study, from our transcriptome, another peanut *CYP707A* gene, *AhCYP707A3* was identified, and AhCYP707A3 protein shared 84.4%, 50.9%, 65%, 54%, 68.2% and 53% sequence identity with AhCYP707A1, 2 (our previously characterized two peanut CYP707As [18]), and AtCYP707A1, 2, 3 and 4, respectively. Like AhCYP707A1 and 2, AhCYP707A3 contains the highly conserved cysteine motif (PFGNGTHSCPG), which was reported to be essential for the hydroxylation [54]. Three peanut CYP707A proteins (AhCYP707A1, 2 and 3) were all predicted as having a signal peptide by the iPSORT algorithm, consistent with the report of ER-membrane localized ABA catabolism catalyzed by CYP707As [20]. In the phylogenetic tree (Fig 5), AhCYP707A1, 3 and AtCYP707A1, 3 proteins were clustered into one group, and AhCYP707A3 was relatively closer to AhCYP707A1, consistent with the above result of sequence identity analysis.

**Fig 5.**
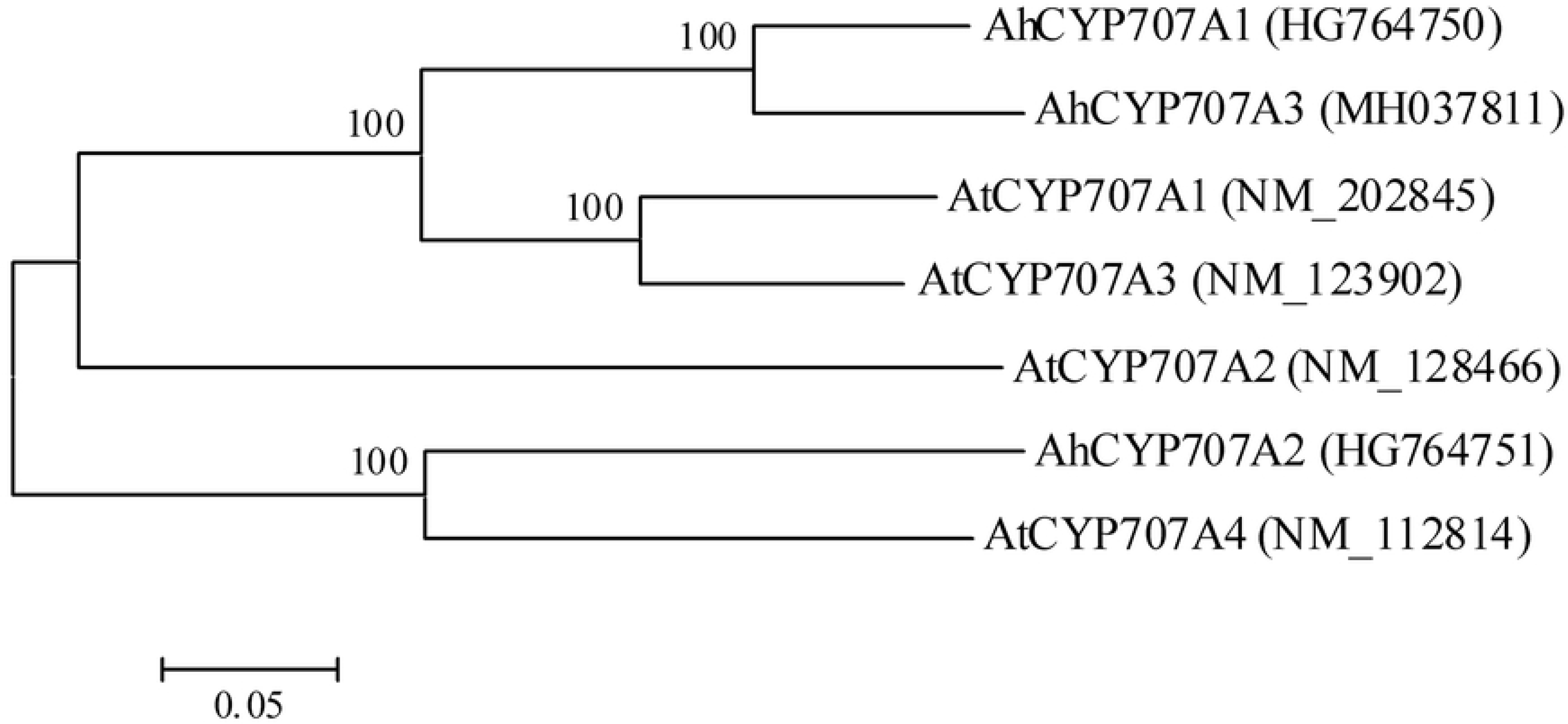
Phylogenetic analysis of amino acid sequences deduced from three peanut (*AhCYP707A1*, *2* and *3*) and four *Arabidopsis* (*AtCYP707A1*, *2*, *3* and *4*) *CYP707A* genes. Multiple sequence alignment was performed using Clustal W and phylogenetic tree was constructed via the Neighbor-Joining method in MEGA 4 software. Bootstrap values from 1000 replicates for each branch were shown. GenBank accession numbers for each aligned CYP707A sequence were indicated in parentheses. The scale bar is 0.05.

The main conjugation pathway for ABA is glucosylation catalyzed by ABA UDP-glucosyltransferases (UGTs), which produces ABA-GE, a storage form and an inactive end product of ABA metabolism [55,56]. Previously reported UGTs, UGT71B6, UGT71B7 and UGT71B8, UGT73B1 and UGT73B3, UGT75B1 and UGT75B2, UGT84B1 and UGT84B2, which displayed *in vitro* the activity to glucosylate ABA, belong to the UGT subfamilies of the family 1 in *Arabidopsis* [57]. In the present study, two unique ABA *UGT* genes, *AhUGT71K1* and *AhUGT73B4*, were identified from the constructed drought-induced transcriptome of peanut leaves. Multiple alignments showed that AhUGT71K1 protein shared the highest sequence identity with *Arabidopsis* UGT71C5, which was very recently confirmed *in vitro* and *in vivo* to play a major role in ABA glucosylation for ABA homeostasis [57]. AhUGT73B4 shared the highest sequence identity with *Arabidopsis* UGT73B1, which displayed ABA glucosylation activity *in vitro* [57]. A motif, named as UDPGT [58], involved in binding to the donor sugar was highly conserved in the C-terminal sequences of all analyzed UGT proteins (Fig 6A). AhUGT71K1 and AhUGT73B4 were both predicted by the WoLF PSORT tool to localize in the cytosol, similar to the cytosolic localization of UGT71B6, UGT71B7, UGT71B8 and UGT71C5 [17,57]. Consistent with the result of sequence alignment, AhUGT71K1 and AhUGT73B4 were phylogenetically closer to UGT71C5 and UGT73B1, respectively (Fig 6B).

**Fig 6.**
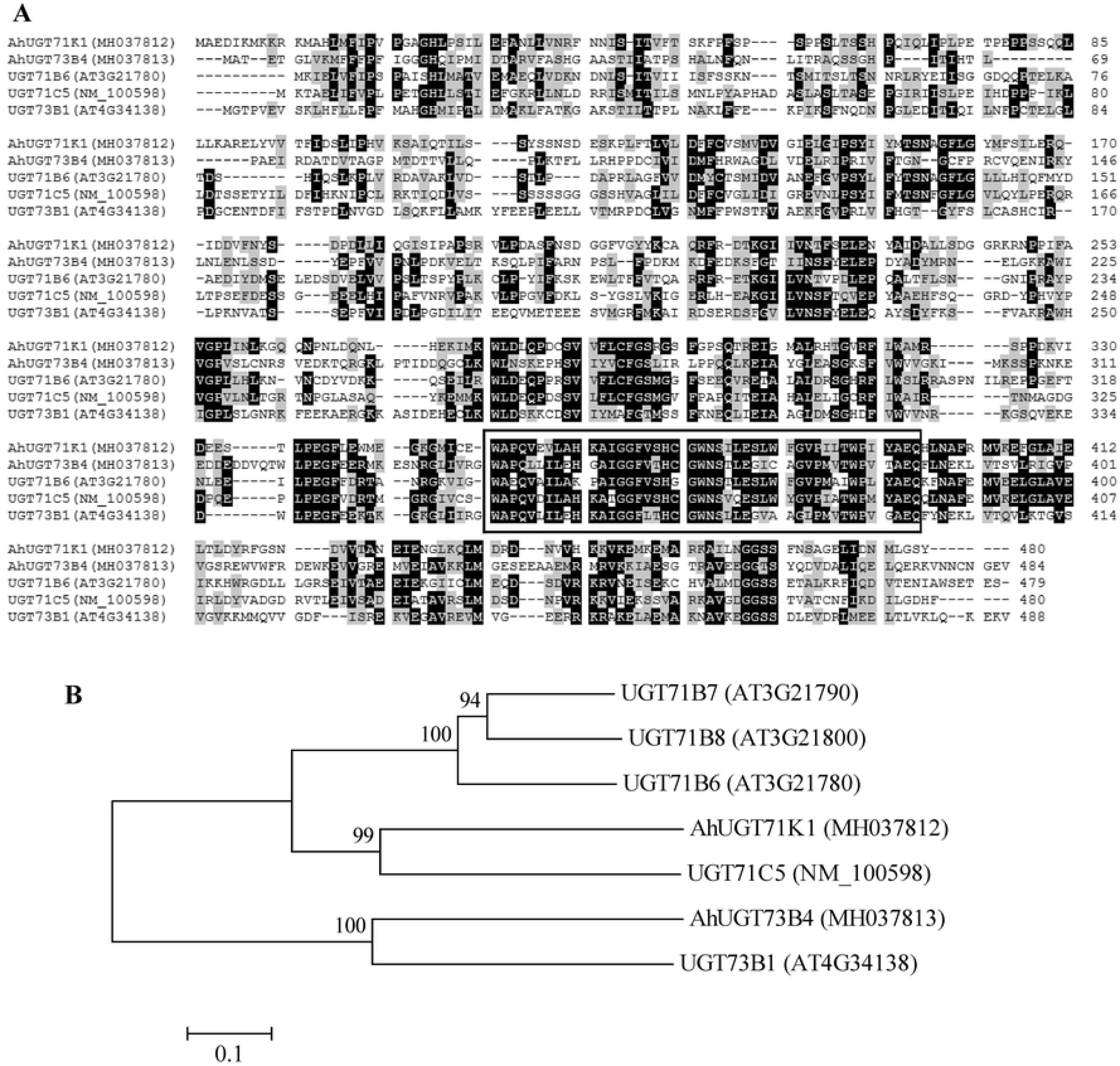
Sequence analyses of UGT proteins from peanut and *Arabidopsis*. **(A)** Alignment of deduced amino acid sequences from peanut *AhUGT71K1*, *AhUGT73B4* and *Arabidopsis UGT71B6*, *UGT71C5*, *UGT73B1*. Identical and similar amino acid residues were shaded in black and gray, respectively. Dotted lines indicated gaps that were introduced to maximize the alignment. The highly conserved motif UDPGT in all UGTs was boxed. Amino acids were numbered from the initial methionine. GenBank accession numbers for each aligned UGT homolog were indicated in parentheses. **(B)** Phylogenetic analysis of amino acid sequences of peanut AhUGT71K1, AhUGT73B4 and *Arabidopsis* UGT71B6, UGT71B7, UGT71B8, UGT71C5, UGT73B1. Multiple sequence alignment was performed using Clustal W and phylogenetic tree was constructed via the Neighbor-Joining method in MEGA 4 software. Bootstrap values from 1000 replicates for each branch were shown. GenBank accession numbers for each analyzed UGT were indicated in parentheses. The scale bar is 0.1.

The translocation of ABA between cells, tissues and organs also plays important roles in whole plant physiological response to stress conditions. ABA can diffuse passively across biological membranes when it is protonated [21,59], and can also be transported across plasma membranes by ABCG transporters [60,61]. To date, at least eight different ABA transporters have been identified by genetic and functional screening [22–29]. In the present study, two *ABCG* gene homologs, *AhABCG22.1* and *AhABCG22.2* were screened and characterized from our constructed drought-induced transcriptome of peanut leaves. Multiple alignments showed that AhABCG22.1 and AhABCG22.2 proteins shared 81% mutual sequence identity; AhABCG22.1 shared 75.6% and 36.7% sequence identity with *Arabidopsis* ABCG22 and ABCG25, respectively; AhABCG22.2 shared 75.3% and 37.8% sequence identity with *Arabidopsis* ABCG22 and ABCG25, respectively. The characterized domains ABC transporter G-25 (residues 111-746 and 123-726 respectively in AhABCG22.1 and AhABCG22.2) and ABC2_membrane (residues 501-703 and 483-685 respectively in AhABCG22.1 and AhABCG22.2) were highly conserved; the conserved features of ATP-binding site, ABC transporter signature motif, Walker A/P-loop and Walker B were also found in both AhABCG22.1 and AhABCG22.2 (Fig 7A). AhABCG22.1 and AhABCG22.2 were both predicted subcellularly as integral plasma membrane proteins. Phylogenetic tree of AhABCG22.1 and AhABCG22.2, and five *Arabidopsis* ABCGs (ABCG25, ABCG40, ABCG22, ABCG30 and ABCG31) demonstrated that AhABCG22.1 and AhABCG22.2 were clustered with ABCG22, and that all three were relatively closer to ABCG25 (Fig 7B).

**Fig 7.**
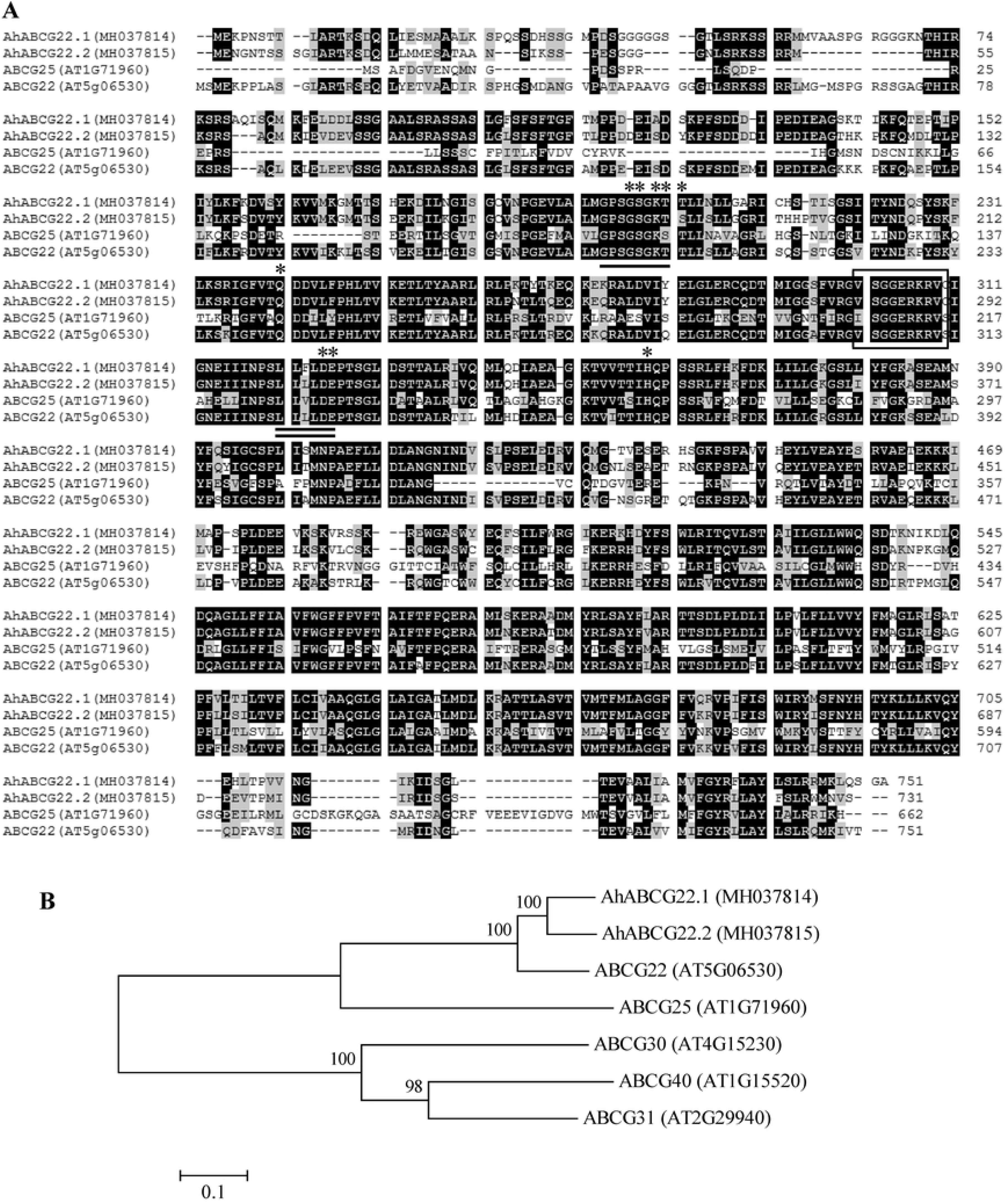
Sequence analyses of ABCG proteins from peanut and *Arabidopsis*. **(A)** Alignment of deduced amino acid sequences from peanut *AhABCG22.1*, *AhABCG22.2* and *Arabidopsis ABCG25*, *ABCG22*. Identical and similar amino acid residues were shaded in black and gray, respectively. Dotted lines indicated gaps that were introduced to maximize the alignment. The highly conserved features of ATP-binding site, ABC transporter signature motif, Walker A/P-loop and Walker B in all ABCGs were respectively indicated with asterisks, a box, an underline and a double-underline. Amino acids were numbered from the initial methionine. GenBank accession numbers for each aligned ABCG homolog were indicated in parentheses. **(B)** Phylogenetic analysis of amino acid sequences of peanut AhABCG22.1, AhABCG22.2 and *Arabidopsis* ABCG25, ABCG40, ABCG22, ABCG30, ABCG31. Multiple sequence alignment was performed using Clustal W and phylogenetic tree was constructed via the Neighbor-Joining method in MEGA 4 software. Bootstrap values from 1000 replicates for each branch were shown. GenBank accession numbers for each analyzed ABCG were indicated in parentheses. The scale bar is 0.1.

### Expression pattern of genes involved in ABA production, catabolism and transport in peanut leaves in response to drought stress

It has been reported that, with the exception of *AtABA2*, the expressions of most of the genes involved in *de novo* biosynthesis of ABA are up-regulated by drought stress [8–12,48,62]. In contrast, *AtABA2* is expressed constitutively at a relatively low level and is not induced by dehydration stress [7,8]. In the present study, real-time RT-PCR was performed to detect the expressions of the above characterized genes involved in ABA biosynthetic pathway in peanut leaves in response to drought stress. The results showed that gene expressions of *AhZEP*, *AhNCED1*, *AhAAO2* and *AhABA3* were all significantly up-regulated in response to drought stress (Fig 8). Particularly, the transcript level of *AhNCED1* gene was strongly up-regulated (756 times higher than that in the control at 10 h of the stress) by drought stress (Fig 8B), consistent with our previous reports [18,34]. The expression of *AhNCED3* (Fig 8D) was also induced (0.9 times higher than that in the control at 10 h of the stress) by drought, but the induction was much slighter than that of *AhNCED1* (Fig 8B, D). However, the expressions of *AhABA2* (Fig 8C) and *AhAAO1* (Fig 8E) were not affected significantly by the stress, which were consistent with the previous reports of *AtABA2* [7,8] and *AhAAO1* [51].

**Fig 8.**
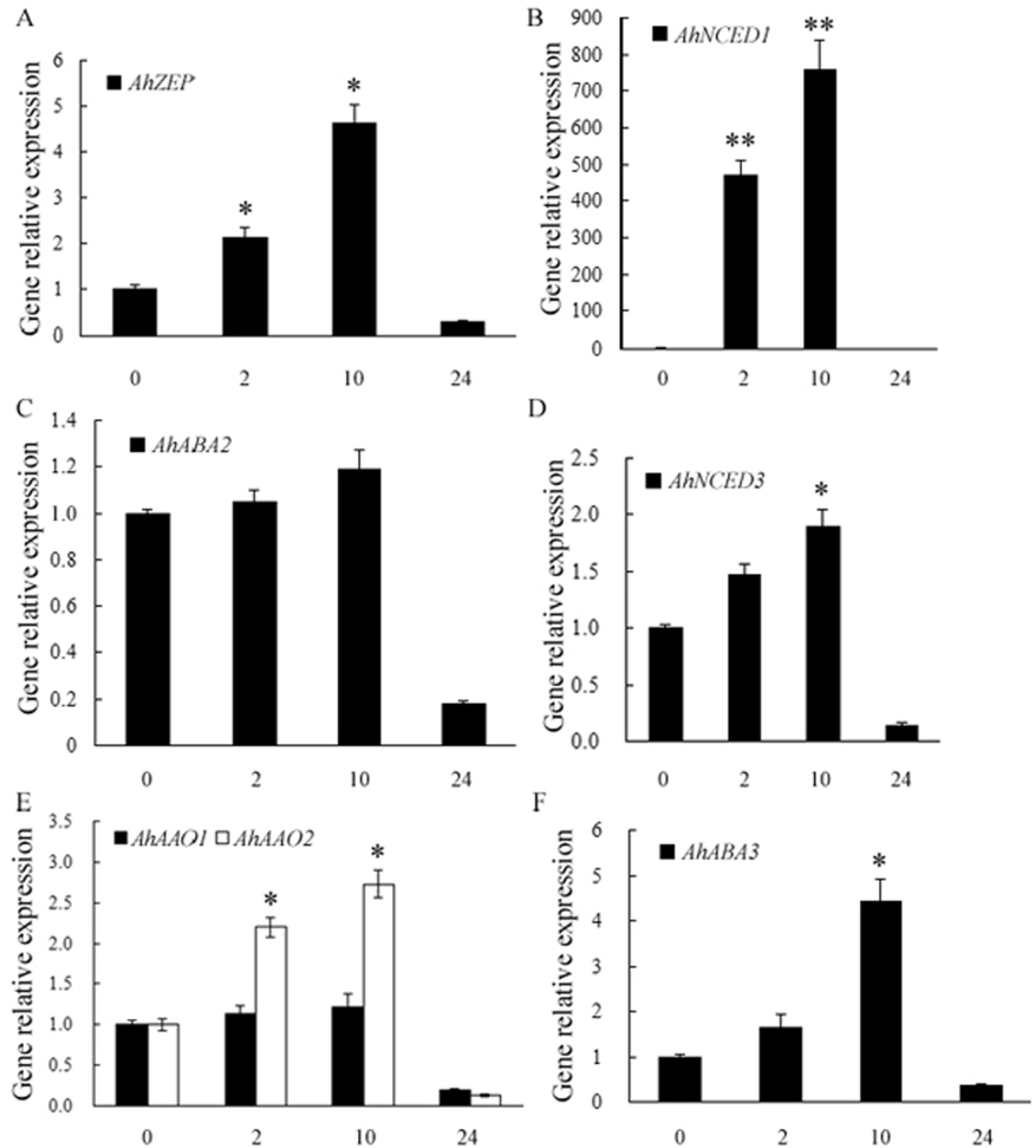
Expressions of ABA biosynthetic genes, including *AhZEP* (A), *AhNCED1* (B) and *AhNCED3* (D), *AhABA2* (C), *AhAAO1* and *AhAAO2* (E), and *AhABA3* (F) in peanut leaves in response to drought stress. Peanut seedlings of twelve days old were hydroponically grown in the solution containing 20% PEG6000 for indicated time, or deionized water as a control (0). Total RNA was prepared respectively from leaves of control or stressed plants. Gene expressions detected by real-time quantitative RT-PCR were shown relative to the expression of peanut *18S rRNA* gene in each sample. All data are presented as mean ± standard errors (SE) of three replicates. The asterisk above each bar indicates a significant difference between stressed and controlled samples at *P* < 0.05 (*) or *P* < 0.01 (**).

Compared with the lengthy *de novo* biosynthetic pathway [3,55,63], the one-step hydrolysis of ABA-GE to ABA catalyzed by BG is a fast process, which is optimal to meet the rapid increase in ABA level in response to stresses. *Arabidopsis AtBG1* and *AtBG2* were both reported to be induced by dehydration stress [13,14]. Loss of *AtBG1* [13] or *AtBG2* [14] in *Arabidopsis* caused lower ABA levels and reduced abiotic stress tolerance, whereas overexpression of *AtBG1* [13] or *AtBG2* [14] resulted in higher ABA accumulation and enhanced tolerance to abiotic stress. In this study, the expressions of *AhBG11* and *AhBG24* genes in peanut leaves in response to drought stress were determined by real-time RT-PCR performance. As shown in Fig 9, the transcript levels of *AhBG11* and *AhBG24* were rapidly and significantly up-regulated by 2-h (4.83- and 4.58-fold increase, respectively) or 10-h (1.97- and 1.65-fold increase, respectively) drought stress.

**Fig 9.**
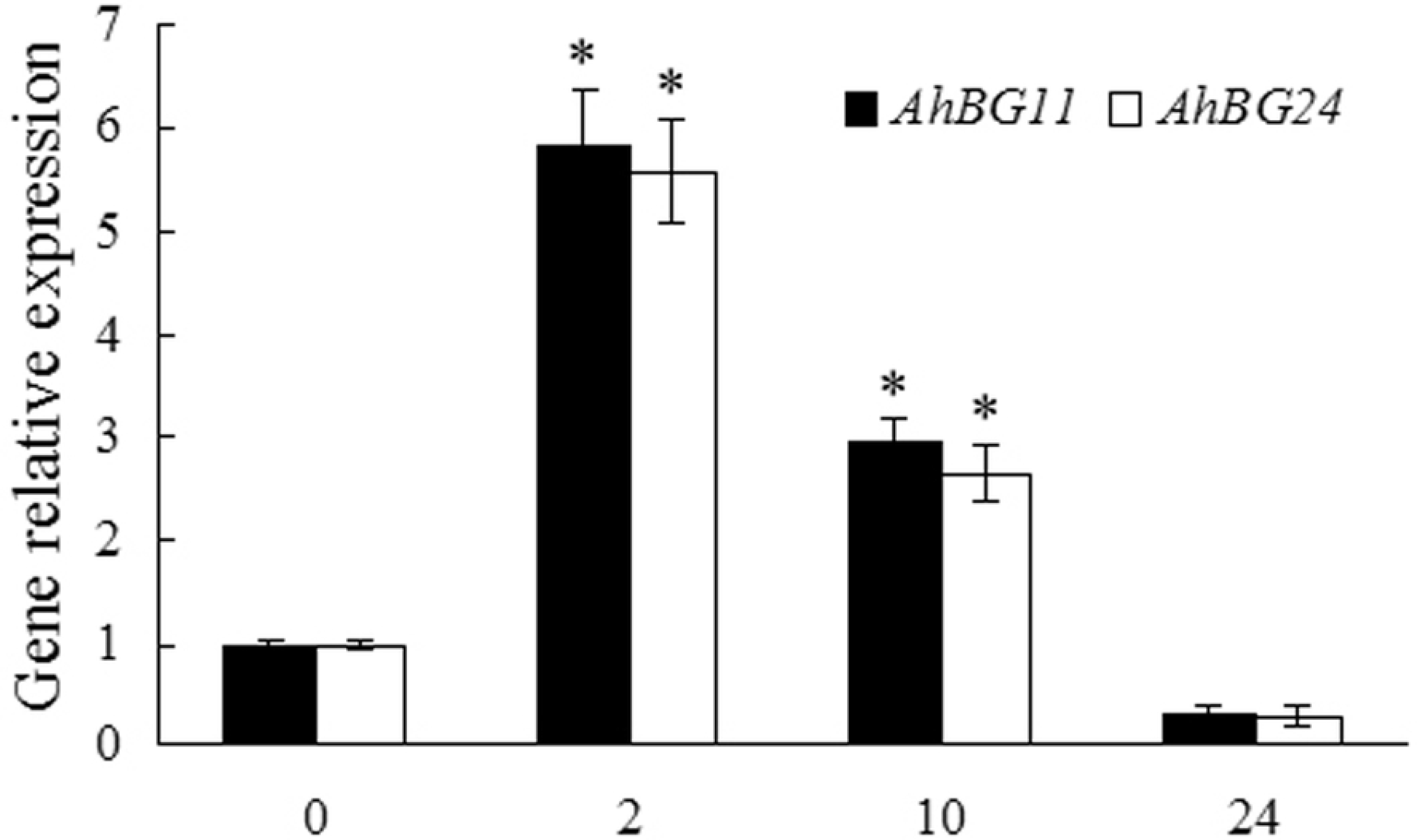
β-glucosidase coding genes, including *AhBG11* and *AhBG24* in peanut leaves rapidly and highly respond to drought stress. Peanut seedlings of twelve days old were hydroponically grown in the solution containing 20% PEG6000 for indicated time, or deionized water as a control (0). Total RNA was prepared respectively from leaves of control or stressed plants. Real-time RT-PCR analysis was performed as described in Fig. 8. All data are presented as mean ± standard errors (SE) of three replicates. The asterisk above each bar indicates a significant difference between stressed and controlled samples at *P* < 0.05 (*).

ABA catabolism is mediated through hydroxylation and glucose conjugation, and also plays important roles in regulating cellular ABA levels. The transcript levels of all four *Arabidopsis CYP707A* genes increased in response to mannitol or drought stress [15]. The *CYP707A5* mRNA level in rice leaves sharply responded to mannitol [64]. We previously demonstrated that the transcript levels of peanut *CYP707A1* and *2* genes increased in response to PEG6000- or NaCl-induced osmotic stress [18]. Here another peanut *CYP707A* gene, *AhCYP707A3* was shown to be significantly induced in leaves in response to drought stress, with a 5.93- or an 8.85-fold increase in the transcript respectively at 2 or 10 h of the stress (Fig 10A). The conjugation of ABA with glucose is catalyzed by UGT to produce ABA-GE [16,17]. In *Arabidopsis*, *UGT71B6* gene and its two homologs, *UGT71B7* and *UGT71B8* were all reported to be rapidly induced by osmotic stress [17]. Liu et al [57] showed that mutation of *UGT71C5* and down-expression of *UGT71C5* in *Arabidopsis* caused delayed seed germination and enhanced drought tolerance; and that overexpression of *UGT71C5* accelerated seed germination and reduced drought tolerance. In the present study, the expression of *AhUGT71K1* gene, highly phylogenetically similar to *UGT71B6* (Fig 6B) was rapidly and significantly increased in peanut leaves in response to drought stress (Fig 10B). Whereas, the transcript level of *AhUGT73B4* in peanut leaves did not respond to drought stress markedly (Fig 10B).

**Fig 10.**
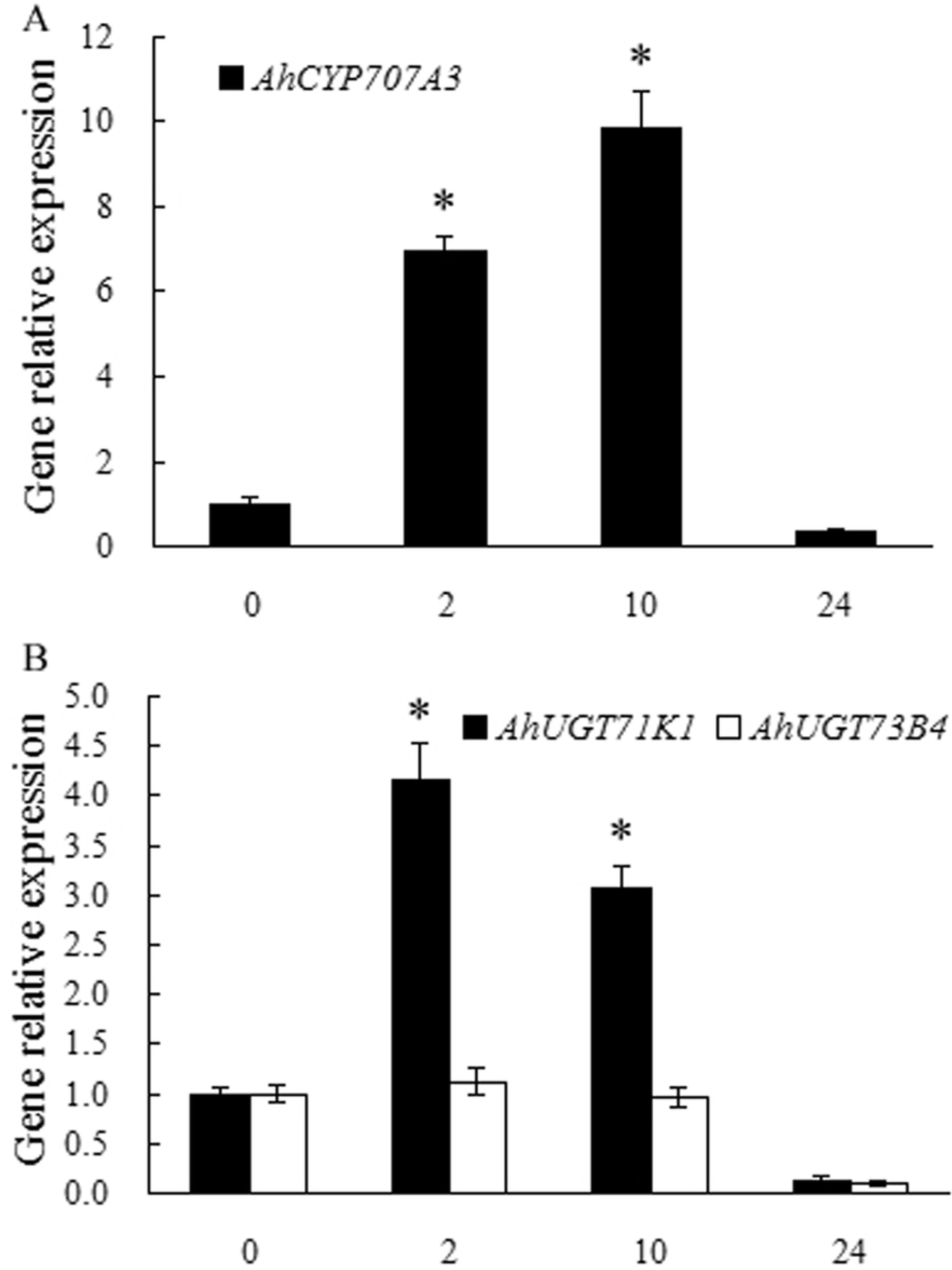
Expressions of ABA catabolic genes, including *AhCYP707A3*, *AhUGT71K1* and *AhUGT73B4* in peanut leaves in response to drought stress. Peanut seedlings of twelve days old were hydroponically grown in the solution containing 20% PEG6000 for indicated time, or deionized water as a control (0). Total RNA was prepared respectively from leaves of control or stressed plants. Real-time RT-PCR analysis was performed as described in Fig. 8. All data are presented as mean ± standard errors (SE) of three replicates. The asterisk above each bar indicates a significant difference between stressed and controlled samples at *P* < 0.05 (*).

*Arabidopsis* ABCG25 and ABCG40 were shown to be responsible for ABA transport and response, which function as an ABA exporter and importer, respectively [22,23]. Recently, the removal of PM-localized ABCG25 via activation of endocytosis and transport to vacuole was confirmed to be another mechanism by which plant cells increase cellular ABA levels in response to abiotic stresses, in addition to the activation of ABA biosynthetic genes [65]. Kuromori et al [24] showed that ABCG22 is required for stomatal regulation and involved in ABA influx. In this study, the expressions of two closely related *ABCG22* genes in peanut leaves, *AhABCG22.1* and *AhABCG22.2*, were significantly up-regulated by 2-h (2.89- and 4.77-fold increase, respectively) or 10-h (1.93- and 2.54-fold increase, respectively) drought stress (Fig 11), respectively. Under abiotic stress conditions, plant cells need to increase the cellular ABA levels to trigger ABA-mediated signaling in order to respond to the stresses [48,66], therefore the expression levels of genes involved in ABA production pathways are up-regulated to increase the cellular ABA levels [8–12,48,62] (Fig 8, 9). At this condition, high levels of *AhABCG22* transcripts would contribute to the rapid increase of cellular ABA levels (Fig 11).

**Fig 11.**
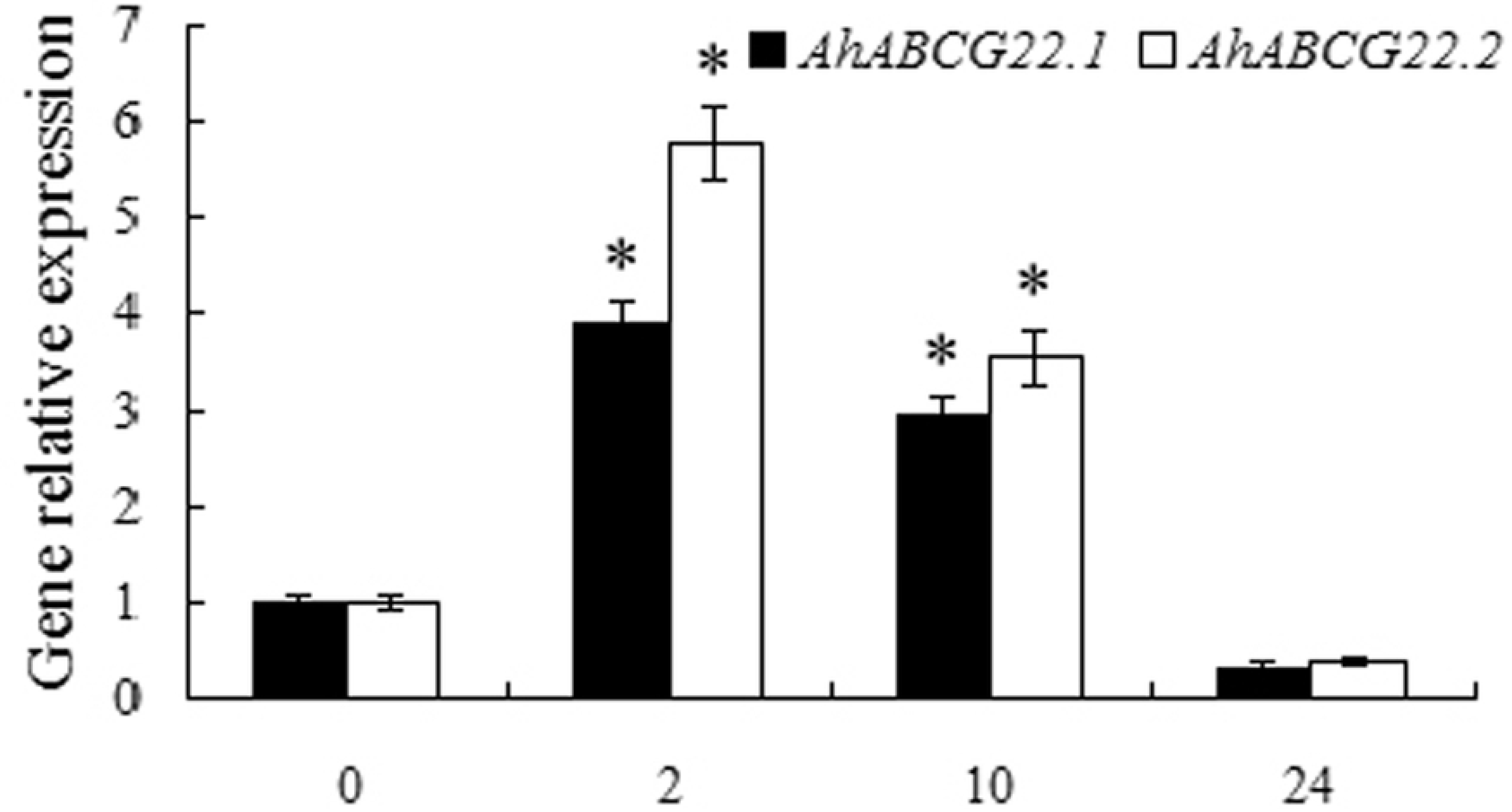
Drought stress significantly induces the expression of ABA importer genes, *AhABCG22.1* and *AhABCG22.2* in peanut leaves. Peanut seedlings of twelve days old were hydroponically grown in the solution containing 20% PEG6000 for indicated time, or deionized water as a control (0). Total RNA was prepared respectively from leaves of control or stressed plants. Real-time RT-PCR analysis was performed as described in Fig. 8. All data are presented as mean ± standard errors (SE) of three replicates. The asterisk above each bar indicates a significant difference between stressed and controlled samples at *P* < 0.05 (*).

### Genes involved in ABA production, catabolism and transport jointly regulate ABA homeostasis in peanut leaves in response to drought

ABA production, catabolism, and transport all affect the active ABA level in plant cells [2]. Two production pathways, *de novo* biosynthesis and hydrolysis of glucose-conjugated ABA, increase the cellular ABA levels [3,13,14,55,63]. The ABA production pathways are activated under abiotic stresses. ABA catabolism via hydroxylation or conjugation to decrease cellular ABA levels is also involved in the regulation of ABA homeostasis [61]. Although extensive work has been performed on the hydroxylation pathway, little is known about the conjugation pathway. In particular, the contribution of conjugation pathway in ABA homeostasis regulation has been less clear. Recently, the determination of ABA content in *Arabidopsis* showed that mutation in *UGT71C5* and down-expression of *UGT71C5* resulted in increased level of ABA, whereas overexpression of *UGT71C5* resulted in reduced level of ABA [57]. The transport of ABA through ABCGs across the plasma membrane is another important pathway to regulate cellular ABA homeostasis [22–24,61]. Consistent with this proposed activity, the ABA exporter *atabcg25* mutants displayed ABA hypersensitive phenotypes at different developmental stages [22]. In contrast, AtABCG40/AtPDR12 is responsible for ABA uptake, which is consistent with the phenotype of *atabcg40*/*atpdr12* that showed a defect in stomatal closure and enhanced water loss [23].

In the present study, the ABA level in peanut leaves in response to 0, 2, 4, 10, 14, 18, or 24 h of drought stress was respectively determined. As shown in Fig 12, the ABA level was significantly increased by drought stress. The ABA content rapidly began to accumulate within 2 h (a 56.6-fold increase) from the start of stress. The highly and rapidly stress up-regulated expressions of genes involved in ABA production and transport, particularly *AhNCED1* (Fig 8B), *AhBG11* and *AhBG24* (Fig 9), and *AhABCG22.1* and *AhABCG22.2* (Fig 11), might contribute to the rapid ABA accumulation (Fig 12).

At 10 h of drought stress, the ABA level reached a peak, 95.9 times higher than that in the control (Fig 12). ABA homeostasis maintained through a balance between the production, catabolism and transport, rather than simply by the biosynthesis. Consistent with this idea, the expressions of genes involved in ABA production (*AhZEP*, *AhNCED1*, *AhABA3*, *AhAAO2*, *AhBG12* and *AhBG24*) (Fig 8, 9) and catabolism (*AhCYP707A3*, *AhUGT71K1*) (Fig 10) were both up-regulated upon drought stress, although the induction of biosynthetic genes (*AhNCED1*) was much higher than that of catabolic genes (*AhCYP707A3* and *AhUGT71K1*). This difference in induction kinetics of gene expression may define the significant accumulation of stress-induced ABA levels (Fig 12). But the ABA content then started to decrease at 18 h of the stress, and reduced to an even lower level than that of the normal (likely due to severe damages induced by drought stress) (Fig 12).

**Fig 12.**
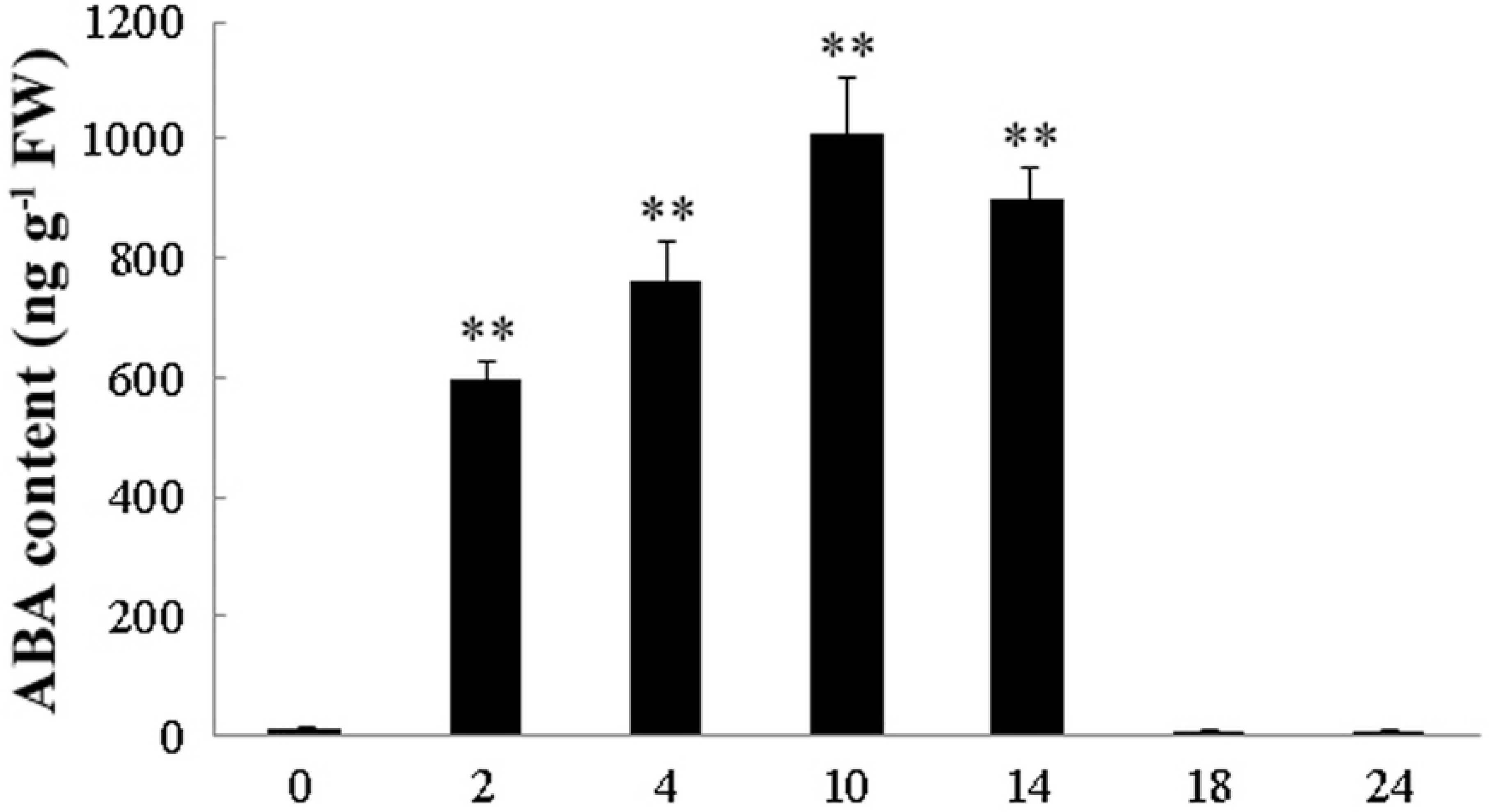
The ABA level in peanut leaves in response to 0, 2, 4, 10, 14, 18, or 24 h of drought stress. The ABA levels in peanut leaves at the presence or absence of drought were determined triplicately for each sample. All data are presented as mean ± standard errors (SE) of three replicates. The asterisk above each bar indicates a significant difference between stressed and controlled samples at *P* < 0.01 (**).

## Conclusions

The two ABA-producing pathways, taking place in different compartments, coordinate to maintain the cellular ABA levels. Additionally, the catabolic pathways play a critical role in the regulation of cellular ABA levels. Furthermore, the PM-localized ABA-specific transporters also contribute to the regulation of cellular ABA levels in plant cells.

The differential subcellular localization of all the key enzymes involved in ABA metabolism and transport indicates that integrated regulatory networks involving multiple organelles are implicated in the regulation of cellular ABA levels. Therefore, a mechanism must exist to achieve a homeostasis of the cellular ABA level that is required for adaptation responses to physiological, developmental, and environmental conditions. However, the entire regulatory network at the molecular level is not fully understood. To elucidate such a mechanism, it is necessary to identify all of the components involved in the regulation of ABA homeostasis, including those that function in production and catabolism, as well as in transport between compartments.

From our previously constructed transcriptome of peanut leaves in response to drought stress, fourteen candidate genes involved in ABA production, catabolism and transport, were identified homologously and phylogenetically (Fig 1-7), and further analyzed at the transcriptional level (Fig 8-11), simultaneously determining ABA level in peanut leaves in response to drought (Fig 12). The high sequence identity and very similar subcellular localization (Table 2) of the proteins deduced from 14 identified genes involved in ABA production (Fig 1-4), catabolism (Fig 5, 6) and transport (Fig 7) with the reported corresponding enzymes in databases suggest their similar roles in regulating cellular ABA levels.

In response to drought stress, ABA accumulation levels in peanut leaves (Fig 12) agree very well with the up-regulated expressions of ABA-producing genes (Fig 8, 9) and PM-localized ABA importer genes (Fig 11), although the expression of ABA catabolic genes was also up-regulated (Fig 10). It is likely that drought-responsive induction of catabolic genes helps not only to maintain ABA levels within a permissible range, but also to prepare the plant for degradation of ABA after removal of the stress. These results suggest that ABA homeostasis in peanut leaves in response to drought may be coordinated by a master regulatory circuit that involves production, catabolism, and as well as transport.

## Author Contributions

**Conceptualization:** Xiaorong Wan, Ling Li

**Data curation:** Haitao Long, Zhao Zheng, Yajun Zhang, Xiaorong Wan

**Formal analysis:** Haitao Long, Zhao Zheng, Yajun Zhang, Pengzhan Xing, Xiaorong Wan

**Funding acquisition:** Pengzhan Xing, Xiaorong Wan, Ling Li

**Investigation:** Haitao Long, Zhao Zheng, Yajun Zhang, Pengzhan Xing, Xiaorong Wan

**Methodology:** Haitao Long, Zhao Zheng, Yajun Zhang, Xiaorong Wan

**Project administration:** Pengzhan Xing, Xiaorong Wan, Ling Li

**Resources:** Xiaorong Wan, Yixiong Zheng, Ling Li

**Software:** Haitao Long, Zhao Zheng, Yajun Zhang, Xiaorong Wan

**Supervision:** Xiaorong Wan, Yixiong Zheng, Ling Li

**Validation:** Xiaorong Wan, Yixiong Zheng, Ling Li

**Visualization:** Haitao Long, Zhao Zheng, Xiaorong Wan

**Writing – original draft:** Haitao Long, Zhao Zheng, Yajun Zhang, Pengzhan Xing

**Writing – review & editing:** Haitao Long, Yajun Zhang, Xiaorong Wan, Ling Li

